# SARS-CoV-2 viability in time on experimental surfaces

**DOI:** 10.1101/2021.03.04.433846

**Authors:** Maria A. Nikiforova, Andrei E. Siniavin, Elena V. Shidlovskaya, Nadezhda A. Kuznetsova, Vladimir A. Gushchin

**Affiliations:** N.F. Gamaleya National Research Center for Epidemiology and Microbiology, Ivanovsky Institute of Virology, Ministry of Health of the Russian Federation, Moscow, Russia; Department of Molecular Neuroimmune Signalling, Shemyakin-Ovchinnikov Institute of Bioorganic Chemistry, Russian Academy of Sciences, Moscow, Russia; Lomonosov Moscow State University, Moscow, Russia

**Keywords:** SARS-CoV-2, viability, surfaces, PCR

## Abstract

We evaluated the SARS-CoV-2 viability preservation on different model surfaces over time. It was found that the SARS-CoV-2 RNA was detected on all studied surfaces for 360 minutes, while the viability of the virus was completely lost after 120 minutes. Type of experimental surface significantly affects viability preservation.

## Text

Environmental surfaces are suspected to be contaminated with the SARS-CoV-2 and are likely sources of COVID-19 transmission (1). The World Health Organization (WHO) has found that there is still not enough scientific evidence of the viability of SARS-CoV-2 on inert surfaces. Scientific reports on the viability of SARS-CoV-2 report that the virus can persist differently according to the surface, from hours to days. For example, the SARS-CoV-2 stable on plastic and stainless steel, copper, cardboard, and glass with durations detected up to 72, 4, 24, and 84 h, respectively (2).

However, the fact that the virus is present on the surface does not mean that the surface itself is dangerous and can become a source of infection (3,4).

Studies show that after a 3-hour incubation the infectious virus is not detected on the paper for printer and napkins or on treated wood and cloth in one day. In contrast, SARS-CoV-2 was more stable on smooth surfaces. Thus 39 non-infectious samples were positive, which indicates that non-infectious viruses could still be detected (5).

Modeling of the SARS-CoV-2 in time viability preservation upon contact with five model materials was carried out in laboratory controlled experimental conditions. The most common materials including ceramic tile, metal (aluminum foil), wood (chipboard), plastic, and cloth (towel) had been used SARS-CoV-2 strain PMVL-3 (GISAID: EPI_ISL_470897) was isolated from naso/oropharyngeal swab and propagated on Vero E6 cells (ATCC CRL-1586). A 15 μl of viral culture the SARS-CoV-2 (containing 0,4*10^5^TCID_50_/ml) was pipetted on a surface (~ 1.5-2 cm^2^) of each material in quintuplicate. Groups of samples material and virus control were incubated for 0 min, 15 min and 30 min (wet surface) or 120 min and 360 min (dried at room temperature). After virus exposure, the virus was eluted from the experimental surface with 200 μL of PBS.

Assessment of the presence of SARS-CoV-2 RNA was carried out by quantitative RT-PCR. Viable virus was determined by tissue culture assay on 293T/ACE2 cells and virus titer was calculated using the Reed and Muench method. The data was processed in the GraphPad Prism 7 software and analyzed using the ANOVA Kruskal-Wallis test. Differences were considered statistically significant at *p* <0.05.

According to the results of the experiments, it was found that SARS-CoV-2 RNA is detected on all experimental surfaces. Significant reduction of **0.5** log_10_ copies/ml SARS-CoV-2 RNA was observed upon contact of the virus with wood (chipboard) for 15 min, as well as on **1** log_10_ copies/ml SARS-CoV-2 RNA on contact with metal and plastic, after 15 min and 30 min respectively. However, in all eluates from experimental materials at both 120- and 360-minutes exposure were detected a high level of SARS-CoV-2 RNA (Figure A). A significant reduction by 1 log_10_ copies/ml of SARS-CoV-2 was noted after exposure for 6 h on a cloth (towel) sample. But, in general, the amount of SARS-CoV-2 RNA was stably high in all kinds of surface and did not differ from virus control (sample not in contact with the material).

**Figure.**
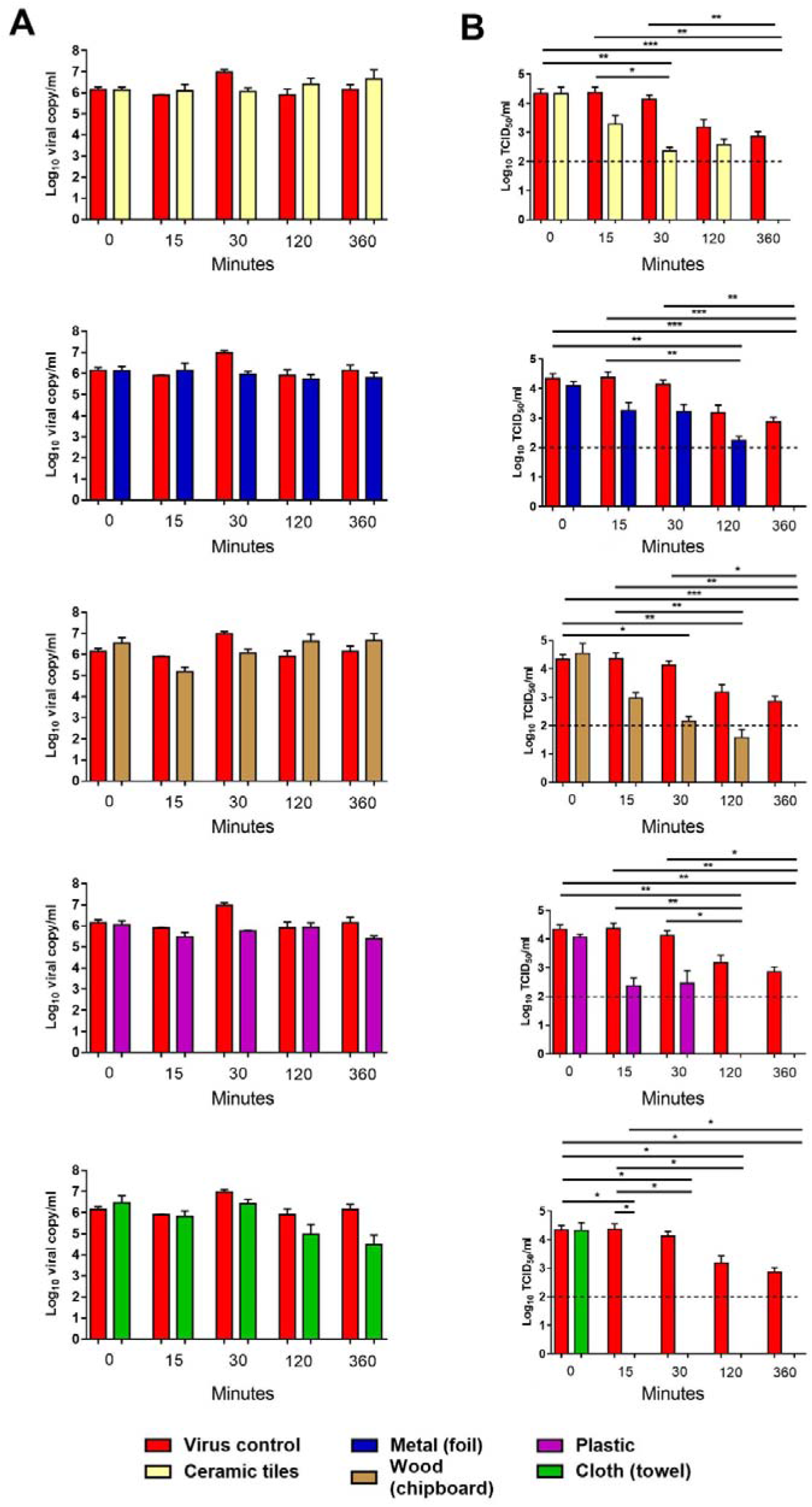
Stability of SARS-CoV-2 on model surfaces under different conditions. Various experimental surfaces were inoculated with 0,4*10^5^TCID_50_/ml SARS-CoV-2 and incubated at room temperature. At indicating time points the virus were eluted and residual virus was detected by A) qRT-PCR or B) viable virus titer was determined by tissue culture assay on 293T/ACE2 cells.

Determination of the infectivity of SARS-CoV-2 after contact with model materials on 293T/ACE2 cells showed a sharp decreased viability of SARS-CoV-2 after 120 min (Figure B). The virus titer gradually decreased depending on the material in the following order: ceramic tile → metal → wood (chipboard) → plastic → cloth (towel). After 120 min exposure of virus on materials such as plastic and a cloth (towel), the infectious virus was not detected while SARS-CoV-2 RNA was still there.

During the assessment of the infectivity of the virus upon contact with model materials it was shown that SARS-CoV-2 RNA is detected on all experimental surfaces, regardless of the conditions and time of exposure to the virus. Even after 360 min the amount of virus on the surface, measured by quantitative RT-PCR, varies insignificant (within the order). However, the detection of SARS-CoV-2 RNA is not indicating the presence of a viable virus. Most significantly reduce the infectivity of the virus when the virus contacts with cloth (towel) samples, as well as plastic. Longer persistence of the infectious virus has been observed on surfaces such as metal, wood (chipboard) and ceramic tile. The decrease in the infectivity of SARS-CoV-2 occurs 120 min after contact with model materials and is completely lost for 360 min of exposure, when drying is achieved. It can be assumed that the complete loss of viability and the infectivity of the virus occurs at an earlier point in time (between 120-360 minutes) for all investigated materials.

Our research is not without its flaws. We used culture fluid to simulate contamination. Its composition will be significantly different from human excreta formed because of natural contact with surfaces. Nevertheless, the results can be useful for planning further research. In the context of environmental safety assessment, the use of RT-PCR alone can lead to highly distorted judgments.

Ms. M. A. Nikiforova is a researcher in the National Research Centre for Epidemiology and Microbiology named after the Honorary Academician N. F. Gamaleya, Moscow. Her primary research interest is the population variability of pathogenic microorganisms.

## Acknowledgments

The authors are grateful to Dr. I.V. Korobko for the general idea and discussion of the study design.

This research was funded by the grant #056 - 00119 - 21-00 provided by the Ministry of Health of the Russian Federation, Russia.

